# sNucConv: A bulk RNA-seq deconvolution method trained on single-nucleus RNA-seq data to estimate cell-type composition of human subcutaneous and visceral adipose tissues

**DOI:** 10.1101/2023.07.16.549187

**Authors:** Gil Sorek, Yulia Haim, Vered Chalifa-Caspi, Or Lazarescu, Maya Ziv, Tobias Hagemann, Pamela Arielle Nono Nankam, Matthias Blüher, Idit F. Liberty, Oleg Dukhno, Ivan Kukeev, Esti Yeger-Lotem, Assaf Rudich, Liron Levin

## Abstract

Deconvolution algorithms rely on single-cell RNA-sequencing (scRNA-seq) data applied onto bulk RNA-sequencing (bulk RNA-seq) to extract information on the cell-types composition and proportions comprising a certain tissue. Adipose tissues’ cellular composition exhibits enormous plasticity in response to weight changes and high variance at different anatomical locations (depots). However, adipocytes – the functionally unique cell type of adipose tissue, are not amenable to scRNA-seq, a challenge recently met by applying single-nucleus RNA-sequencing (snRNA-seq). Here we aimed to develop a deconvolution method to estimate the cellular composition of human visceral and subcutaneous adipose tissues (hVAT and hSAT, respectively) using snRNA-seq to assess the true cell-type proportions. To correlate deconvolution-estimated cell-type proportions to true (snRNA-seq -derived) proportions, we analyzed seven hVAT and 5 hSAT samples by both bulk RNA-seq and snRNA-seq. snRNA-seq uncovered 15 distinct cell types in hVAT and 13 in hSAT. Deconvolution tools – SCDC, MuSiC, and Scaden exhibited low performance in estimating cell-type proportions (median |R|= 0.12 for estimated vs. true correlations). Notably, estimation accuracy somewhat improved by decreasing the number of cell-types groups, which nevertheless remained low (|R|<0.42). We therefore developed **sNuConv**, a novel method that employs Scaden, a deep-learning tool, trained using snRNA-seq - based data corrected by i. snRNA-seq/bulk RNA-seq highly-correlated genes, ii. corrected estimated cell-type proportions based on individual cell-type regression models. Applying sNuConv on our bulk RNA-seq data resulted in cell-type proportion estimation accuracy with median R=0.93 (range:0.76–0.97) for hVAT, and median R=0.95 (range:0.92–0.98) for hSAT. The resulting model was depot-specific, reflecting depot-differences in gene expression patterns. Thus, we present sNuConv, a novel, AI-based, method to deduce the cellular landscape of hVAT and hSAT from bulk RNA-seq data, providing proof-of-concept for producing validated deconvolution algorithms for tissues un-amenable to single-cell RNA sequencing.

## 1. Introduction

Two seminal papers in 2003 demonstrated that obesity is associated with increased adipose tissue abundance of macrophages, whose presence associates with metabolic dysfunction (insulin resistance) (1, 2). These papers sparked enormous interest in deciphering the dynamic cell-composition of adipose tissue. Indeed, over the following 20 years, most known immune cell types had been identified in adipose tissue and were implicated in linking adipose tissue changes with health risks (3, 4). For example, it became clear that it is not only adipose tissue mass per-se, but its cellular composition importantly differentiates people with obesity with cardiometabolic complications from those with a relatively benign obesity phenotype (arguably termed “healthy obesity”) (5). Thus, to understand adipose tissue biology, function, and its interphase with whole-body (patho)physiology, the tissue’s cellular composition must be assessed (6, 7).

With the advent of single-cell RNA-sequencing (scRNA-seq) technology, novel sub-types of specific cells could be studied in an unprecedentedly unbiased approach. However, adipose tissue poses a unique challenge when trying to apply scRNA-seq: the “adipose cells” – adipocytes - that are the unique functional cell type of this tissue, could not be captured by scRNA-seq. This is because adipocytes are unique in several aspects: they have an exceptionally broad size range, which can vary by >20-fold; they are buoyant and also fragile (due to their high lipid content – 85% of the cell volume is composed of triglycerides). Thus, most scRNA-seq platforms, which rely on either microfluidics or the passage of cells through nozzles, disintegrate these cells, particularly those in the larger size range. scRNA-seq approaches have therefore been confined to analyses only of the non-adipocytes, stromal-vascular cell fraction of the tissue. Indeed, such studies uncovered novel cell subtypes that comprise adipose tissue, underscoring the promise of analyzing adipose tissue at the single-cell level.

More recently, single-nucleus RNA-sequencing (snRNA-seq) was developed (8, 9) and applied to studying adipose tissue (10, 11). snRNA-seq’s major weakness in comparison to scRNA-seq is that the nuclear mRNA repertoire is a fraction of the entire cellular RNA landscape. The latter is dominated by cytoplasmic RNA, while nuclear RNA has higher proportions of “immature” mRNA (i.e., that has more intronic regions) and long non-coding RNAs (8, 12). Nevertheless, several studies have demonstrated good correspondence in capturing the transcriptome landscape using scRNA-seq and snRNA-seq (8, 12, 13). On the other hand, snRNA-seq advantages over scRNA-seq have also been noted, in particular, the ability to use flash-frozen (and even formaldehyde-fixed) samples, avoiding the possible transcriptome changes occurring during tissue disintegration into isolated cells. An atlas of human and mouse adipose tissues’ cellular landscape based on snRNA-seq data was recently reported, highlighting known adipose tissue cell types and novel, unbiasedly-identified, sub-types of adipocytes, endothelial cells, and progenitor cells (10).

Despite the advances offered by both scRNA-seq and snRNA-seq, their use remains limited due to their high cost and heavy reliance on specific equipment, reagents, and bioinformatics expertise. Deconvolution tools were therefore developed using computational approaches to address these shortcomings, allowing to estimate cell-type composition of tissues from bulk-tissue RNA-sequencing data. Yet, most algorithms rely on the gene expression derived from scRNA-seq data. Consequently, the ability of such deconvolution tools to accurately estimate adipose tissue’s cell composition, whose true cell-type proportions can only be studied using snRNA-seq gene, is questionable.

Given the importance of characterizing adipose tissue cellular composition, we generated a dataset comprising of human subcutaneous and visceral adipose tissues samples (hSAT and hVAT, respectively), each analyzed in parallel by bulk RNA-seq and snRNA-seq. We tested the ability to infer cell-type composition from bulk RNAseq using available deconvolution tools and modified an existing deep-learning deconvolution algorithm in order to generate a validated deconvolution method for human VAT and SAT, effectively bridging the gap between cytoplasmic and nuclear RNA repertoires.

## 2. Methods

### 2.1 Human adipose tissue samples

Seven hVAT and 5 hSAT samples were obtained from an adipose tissue bio-bank established in Beer-Sheva, Israel, and subjected to both bulk RNA-seq and snRNA-seq. In the Beer-Sheva human adipose tissue biobank, adipose tissue biopsies are obtained from patients who had signed in advance a written informed consent before undergoing elective abdominal surgeries, and tissues are immediately delivered from the operating room to the lab, where they are frozen at -80 C, as detailed previously (14, 15). Ethical approval of the study procedures was obtained by the Helsinki Ethics Committee of Soroka University Medical Center (approval no: 15-0348). Samples included in the present analyses are from consecutive patients with at least 1gr biopsy available, and whose individual basic clinical characteristics are shown in **Supplemental Table 1**.

### 2.1 Bulk RNA Sequencing

Total RNA was isolated from hSAT and hVAT samples (150-250 mg) using RNeasy Lipid Tissue Mini Kit (QIAGEN, Germany), according to the manufacturer’s instructions. RNA quality (RIN^e^) was determined by 4150 TapeStation System (Agilent, CA, USA) with RNA ScreenTape (Agilent, CA, USA), while RNA concentration was determined using Qubit 4 Fluorometer (ThermoFisher Scientific, MA, USA) using Qubit RNA Assay Kit. Only samples with RIN^e^>8.0 were further processed. RNA libraries were prepared using KAPA Stranded RNA-seq Kit (Roche, Switzerland), pooled, and subjected to 76 bp paired-end sequencing according to the manufacturer’s protocol (NovaSeq 6000, Illumina, CA, USA). An average of 47.8 million paired-reads per sample were generated. Sequences were quality trimmed and filtered using Trim Galore (v0.4.4) and cutadapt (v1.15). Alignment of the reads to the human genome (GRCh38.100) was performed with STAR (v2.5.2a) (16). The number of reads per gene per sample was counted using RSEM (v1.2.28) (17). Quality assessment of the process was carried out with FastQC (v0.11.8) and MultiQC (v1.0.dev0) (18). After trimming each sample had an average of 40.5M ± 7.8M reads with an average sequence length of 74bp.

### 2.2 Single-nucleus RNA Sequencing

Human adipose tissue samples were analyzed by isolating nuclei, barcoding them using Chromium 10X technology, followed by RNA sequencing using NovaSeq 6000 (Illumina, CA, USA). Briefly, nuclei were isolated from 500 mg of frozen hSAT or hVAT samples with all steps carried out on ice. One ml of ice-cold adipose tissue nuclei lysis buffer (AST: 5 mM PIPES, 80 mM KCl, 10 mM NaCl, 3 mM MgCl_2_ and 0.1% IGEPAL CA-630) supplemented with 0.2U/µl of RNAse Protector (Roche, Switzerland), were added to each sample, followed by mincing the frozen tissue using small surgical scissors. Then, small pieces of minced tissue in 1 ml of AST lysis buffer were transferred into 7 ml WHEATON® Dounce Tissue Grinder (DWK Life Sciences, Germany) and additional 2 ml of AST were added. Tissues were dissociated to retrieve nuclei by 12 strokes with loose pestle (A), followed by 10 strokes with tight pestle (B). Samples were then incubated for 7 minutes on ice to ensure maximal nuclei retrieval, filtered through 70 µm SMARTStrainer (Miltenyi Biotech, Germany), and centrifuged (4 C, 5 minutes, 500 RCF). Supernatants were discarded, and nuclei pellet was resuspended in 2.5 ml of ice-cold PBS 0.5% BSA, supplemented with 0.2U/µl of RNAse Protector, and filtered through 40 µm SmartStrainer, followed by additional centrifugation. Next, 2 ml of supernatant were discarded, and nuclei pellet was resuspended in the remained 0.5 ml of ice-cold PBS 0.5% BSA. Nuclei were counted using LUNA-FL™ Dual Fluorescence Cell Counter (logos biosystems, South Korea), and 13,000-16,500 nuclei/sample were immediately loaded onto 10X Chromium Controller (10X GENOMICS, CA, USA) according to the manufacturer’s protocol (Chromium Next GEM Chip C. Chromium Next GEM Single Cell 3’ Kits v3.1). cDNA and gene expression libraries size fragments were assessed using 4150 TapeStation System (Agilent, CA, USA) with high sensitivity D5000 ScreenTape, while their concentrations were quantified using Qubit 4 Fluorometer using Qubit dsDNA HS Assay Kit (ThermoFisher Scientific, MA, USA). Gene expression libraries were sequenced by NovaSeq 6000 using S1-100 flow cell (Illumina, CA, USA).

### 2.3 Quality control and mapping

Reads were transformed into a raw counts matrix with CellRanger software, which was used as input for CellBender to remove ambient RNA. For downstream analysis, only cell barcodes which were determined to be true cells by both CellRanger and CellBender algorithms were used (with default parameters). The result of CellBender’s counts matrix (including only filtered barcodes) of each sample was analyzed with Seurat v3.0 R package. Our quality control procedure included removal of broken nuclei, i.e., nuclei with less than 200 genes or nuclei that contained more than 20% mitochondrial genes. Parameters of each sample are presented in **Supplemental Table 2**.

### 2.4 Integration and clustering

Counts were normalized (Log-normalization) and scaled using the 3,000 most variable genes. Linear dimensionality reduction via principal component analysis (PCA) and graph-based clustering via k-nearest-neighbors (KNN) were used to group individual cells into subsets. We removed cell clusters with low mean gene count and low expression level (mean read counts less than one standard deviation below the sample’s mean). We then used the DoubletFinder R package to remove possible doublets. Once each sample passed quality control, we integrated separately the 7 hVAT and 5 hSAT using Harmony and performed dimensionality reduction and clustering again to obtain the final Uniform Manifold Approximation and Projection (UMAP) plots (**Figure1B,C**, respectively).

**Figure 1:**
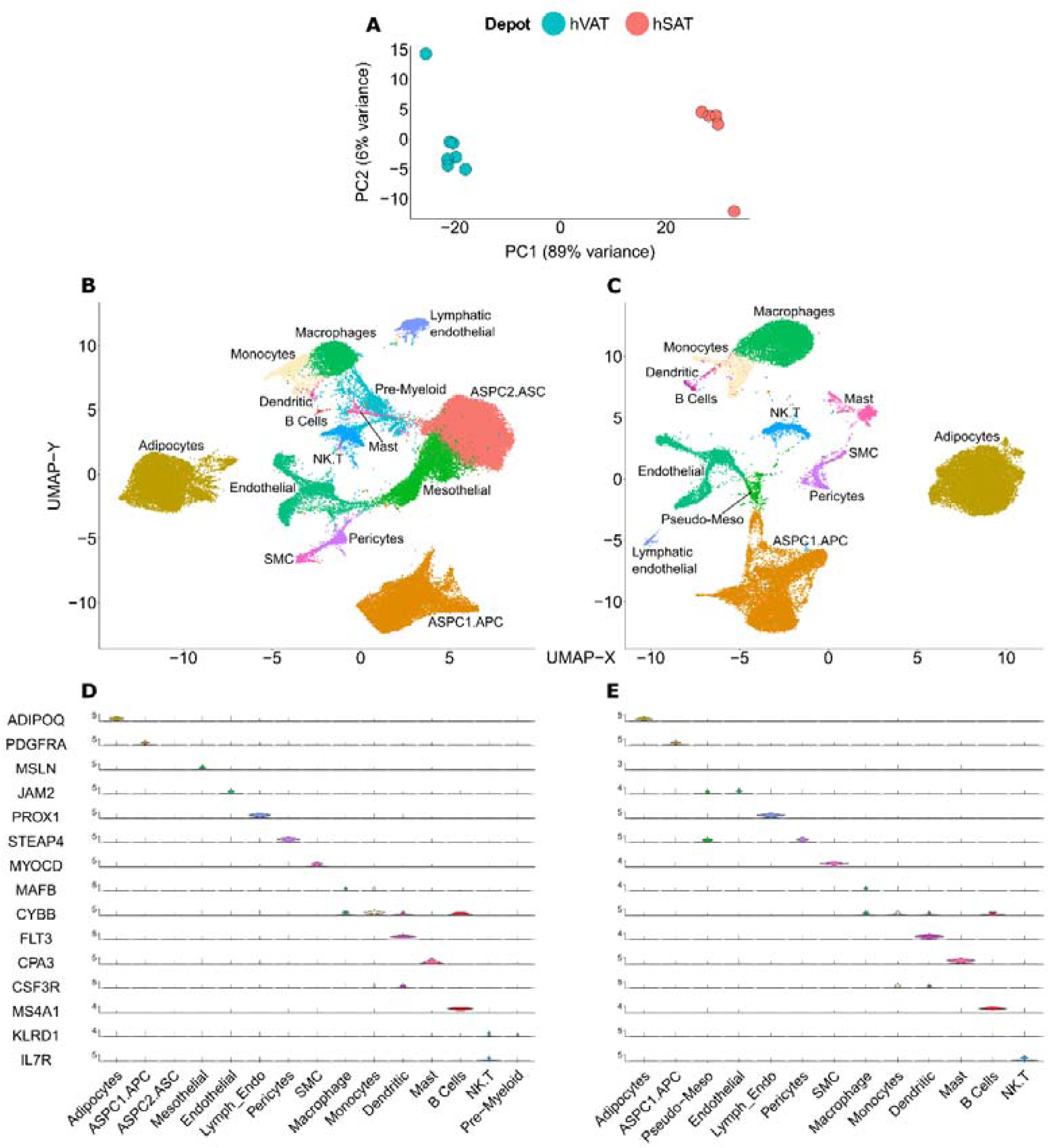
Bulk RNA-seq and single-nucleus RNA-sequencing (snRNA-seq) analysis of human adipose tissues. **A.** Principal Component Analysis (PCA) of bulk RNA-seq analyses of human visceral and subcutaneous adipose tissues (hVAT and hSAT, n=7 and 5 samples, respectively). **B-E.** snRNA-seq analyses of the same hVAT and hSAT as in A, (64,729 and 37,357 nuclei, respectively), performed using 10x genomics platform. UMAP projection uncovering 15 and 13 uniquely colored annotated clusters of nuclei based on analyses of nuclear transcriptome of hVAT and hSAT (**B.,C.**, respectively). Cell-type annotation of nuclei clusters based on single marker genes, as recently used in (10), in hVAT and hSAT (**D., E.**, respectively). ASPC1.APC – adipose stem progenitor cells 1. Adipocyte progenitor cells; ASPC2.ASC - adipose stem progenitor cells 2. Adipose-derived stem cells; SMC – smooth muscle cells

### 2.4 hVAT snRNA-seq annotations

snRNA-seq analysis of the 7 hVAT samples included, after quality control (QC), the nuclear transcriptome of 64,729 nuclei (an average of 9,247 nuclei per sample). UMAP projection map unbiasedly identified 15 unique clusters (**Figure 1B, Supplementary Figure 1A**). Our initial cluster/cell-type annotation relied on single cell-type specific gene markers used in an atlas of the human adipose tissue cellular landscape (10)(**Figure 1D**). Adipocytes, adipocyte progenitor/stem cells (ASPC), mesothelial cells, endothelial cells, lymphatic endothelial cells, pericytes, smooth muscle cells (SMC), mast cells, and B-lymphocytes were easily identifiable using these single cell-type specific markers. Myeloid cells (macrophages, monocytes and dendritic cells) shared more than one marker, and lymphoid T and NK cells clustered into a single cluster. In addition to these readily identifiable clusters, two clusters were largely negative to all 15 markers (see further discussion on their annotation below). To support our clustering, we also searched unbiasedly the 10 highest preferentially (compared to all other clusters) expressed genes (heatmap - **Supplementary Figure 1C,** list of all 10 genes/cluster **- Supplemental Table 3**). This analysis confirmed that indeed the identified clusters exhibited at least 10 differentially-expressed genes per cluster when compared to all other clusters. Ten genes were mildly upregulated in one of the two clusters that were negative to all single marker genes used in **Supplementary Figure 1B**. Expression levels of these genes were also similarly observed in an adjacent cluster, which was identified as mesothelial cells based on an additional set of 10 genes and expression of Mesothelin (MSLN, **Supplementary Figure 1B**). We therefore named this cluster ASPC2.ASC, representing PDGFRA-negative adipocyte stem cells. The myeloid cell clusters overlapped in their up-regulated genes, and a final non-annotated cluster based on single gene markers seemed to express relatively low levels and/or percentage of myeloid markers nuclear RNA, and was therefore named “pre-myeloid”. All samples contributed to each of the cell-type clusters, except for B-cells, to which samples v3387 and v3399 contributed 0 nuclei.

### 2.5 hSAT snRNA-seq annotation

snRNA-seq analysis of the 5 hSAT samples included the nuclear transcriptome of 37,357 nuclei (an average of 7471 nuclei/sample). UMAP projection map unbiasedly identified 13 unique clusters (**Figure 1C**). Single cell-type specific marker genes, depicting the same fifteen single marker genes that were used for the hVAT annotation analysis, are shown in a violin plot (**Figure 1E**). The main 12 cell-type groups (adipocytes, ASPC, endothelial, lymphatic endothelial, pericytes, SMC, mast, B-lymphocytes, macrophages, monocytes, dendritic, lymphoid T and NK cells) that were identified in the hVAT analysis, were also easily identifiable in the hSAT. As reported previously (10), nuclei of cells annotated as mesothelial cells based on the expression of mesothelin were absent in hSAT. In addition, nuclei of cells annotated as PDGFRA-negative adipocyte stem cells (ASPC2.ASC) in the hVAT were also absent in the hSAT. Beyond these readily identifiable clusters, one unique cluster showed co-expression of the endothelial and pericytes marker genes (see further discussion on his annotation below). To support our clustering, we again identified at least 10 preferentially expressed genes in each cluster (heatmap – **Supplementary Figure 2C**, list of all 10 genes/cluster - **Supplemental Table 4**). The previously mentioned unique cluster again overlapped with the top 10 genes for endothelial and pericytes. However, it also exhibited at least 10 unique markers, confirming that it was not a cluster of doublets composed of two nuclei – of endothelial and pericytes. Intriguingly, these same preferentially-expressed genes were also present within the mesothelial cells cluster identified in the hVAT cohort analysis, suggesting that these cells are mesothelial-like, and the cluster was therefore named “Pseudo-Meso”. All samples contributed to each of the cell-type clusters, except for B-cells, to which samples s3387 and s3399, similarly to their corresponding VAT samples, contributed 0 nuclei.

### 2.8 Deconvolution tool assessment

We first assessed three deconvolution tools (Scaden, MuSiC and SCDC) using a leave-one-out approach. Specifically, for each tool, one sample at a time was used as a test sample, while all other samples were used for training. In each run, the test sample’s estimated proportions by each of the tools were compared to the true, snRNA-seq -derived, proportions, and R Pearson coefficient was used to quantify the correlation accuracy. MuSiC requires raw read counts for both bulk RNA-seq and single-cell expression (snRNA-seq, in our case). The prediction was done according to the ’estimation of cell type proportions’ protocol in MuSiC’s documentation. Data preparation included the creation of bulk RNA-seq and snRNA-seq ExpressionSets (also known as ESETs), holding the expression data along with sample and feature annotation. Cell-type proportions were estimated with the ’music_prop’ function, while using all marker genes that were overlapping in the bulk and snRNA-seq ESETs. All other arguments were set to default. SCDC takes ExpressionSet objects with raw read counts as input. The prediction protocol was performed according to SCDC’s vignettes. The single-cell (snRNA-seq, in our case) ESET was first pre-processed, and a quality control procedure was performed to remove cells with questionable cell-type assignments. The processed ESET was evaluated using the ’SCDC_qc’ function, with a threshold set to 0.7. The cell-type proportions were then estimated using the ’SCDC_prop_ONE’ function and default settings, with the processed ESET used as a reference. Scaden requires two input files: (i) cell-type labels file of size (n x 1), where n is the number of cells in the data, and a single ’Celltype’ column with cell-type assignments; (ii) count data file of size (n x g), where g is the number of genes and n is the number of cells (nuclei, in our case). These files were then used to create pseudo bulk samples for training using the ’simulate’ step, generating 1000 artificial samples from 1000 cells per sample. Scaden was then used to process the training data using the ’process’ step, creating a new file for training, which only contains the intersection of genes between the training and the estimation data. The variance cutoff used for the process step was 0.1. Scaden was then trained on three deep neural network models using the ’train’ step, each of them trained for 5000 steps. As a final step, the cell-type proportions were predicted using the ’predict’ step, by providing Scaden bulk count data with gene counts over samples.

### 2.9 Statistical analyses

Correlation coefficients and statistical significance were performed with R (v4.1.1). A p-value <0.05 was considered statistically significant. The development of a new deconvolution algorithm (sNuConv) is described in results section 3.2.

## 3. Results

To develop a deconvolution tool that would reliably estimate cell-type composition of human adipose tissue from bulk RNA-seq, we generated a dataset consisting of 7 human visceral adipose tissue (hVAT) samples and 5 human subcutaneous adipose tissue (hSAT) samples (**Illustrated workflow is presented in Figure 2**). Each of the 12 adipose tissue biopsies was analyzed in parallel by both snRNA-seq and bulk RNA-seq (**Figure 2A**). Patients’ basic characteristics, and Cell Ranger -based quality control (QC) parameters of each sample are shown in **Supplemental Tables 1 and 2**, respectively.

**Figure 2:**
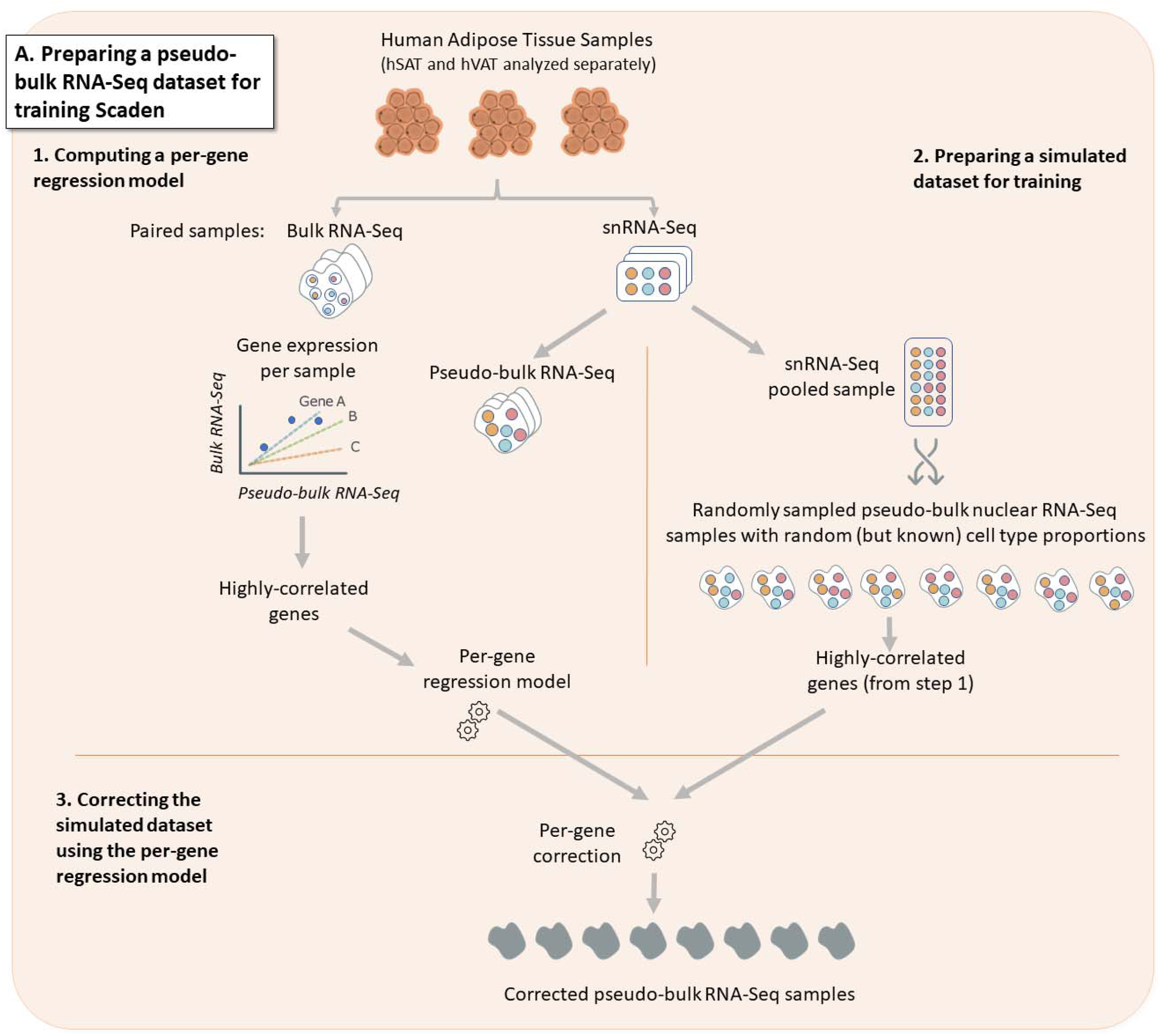
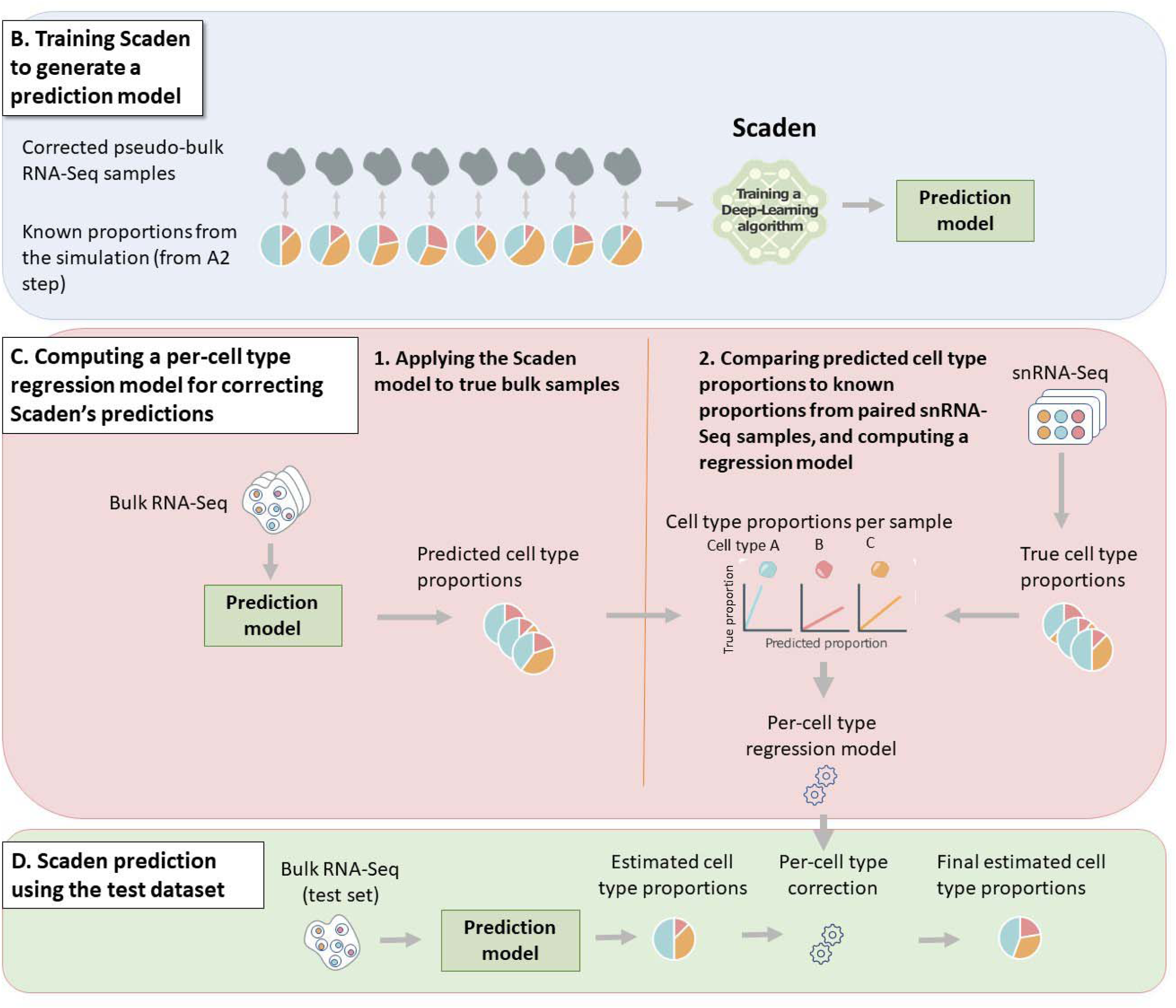
sNuConv workflow. sNuConv is a Scaden-based algorithm developed for estimating cell-type proportions from bulk RNA-seq data while training on snRNA-seq, rather than scRNA-seq, data. **A-D** denote the 4 stages of sNuConv, which include the conversion of snRNA-seq dataset into pseudo-bulk training set with per-gene correction (**A.**), generating a Scaden-based prediction model (**B.**), per cell-type regression model generation (**C.**) and correction (**D.**) to obtain the final sNuConv deconvolution output.

Recent snRNA-seq studies consolidated (harmonized) the sequencing results obtained from the two adipose tissue (anatomically-distinct) depots - hVAT and hSAT (10). Yet, these two adipose tissue depots express depot-specific genes and differ in their gene expression profiles (19–21). Consistently, bulk RNA-seq principal component analysis (PCA) reveals that the global transcriptome of hSAT and hVAT are in distinctly separable clusters **Figure 1A**. Moreover, a multitude of studies demonstrate differences in the function and contribution of the two tissue depots to whole-body physiology (reviewed in (22, 23)). We therefore considered the various deconvolution tools separately for hVAT and hSAT.

### 3.1. Performance of existing deconvolution tools applied onto bulk RNA-seq of human adipose tissue using snRNA-seq data

In order to assess the performance of existing deconvolution tools in uncovering cell-type proportions from bulk RNA-seq of hSAT and hVAT, we took advantage of our unique paired dataset of snRNA-seq and bulk RNA-seq. We tested three tools, two of which utilize cell-type specific gene expression: MuSiC (which was also used in (10)) and SCDC. The third tool – Scaden, uses deep neural network ensemble training. We compared the accuracy of each of the three tools, as detailed in Methods, using a leave-one-out approach: one sample at a time was used as a test sample, while all other samples were used for training. The tools’ accuracy was assessed by evaluating the median R-correlation coefficient between the true, snRNA-seq-derived cell types’ proportions (**Figure 3A, 3C** for hVAT and hSAT, respectively), and the proportions estimated by each deconvolution tool. The three deconvolution algorithms estimated the relative proportions of the 15 hVAT and 13 hSAT different cell types with low accuracy, indicated by a median correlation coefficient of |R|=0.1 (**Figure 3B and 3D**, respectively). Interestingly, all tools exhibited both positive and negative correlations between true and estimated cell-type proportions, particularly in hVAT.

**Figure 3:**
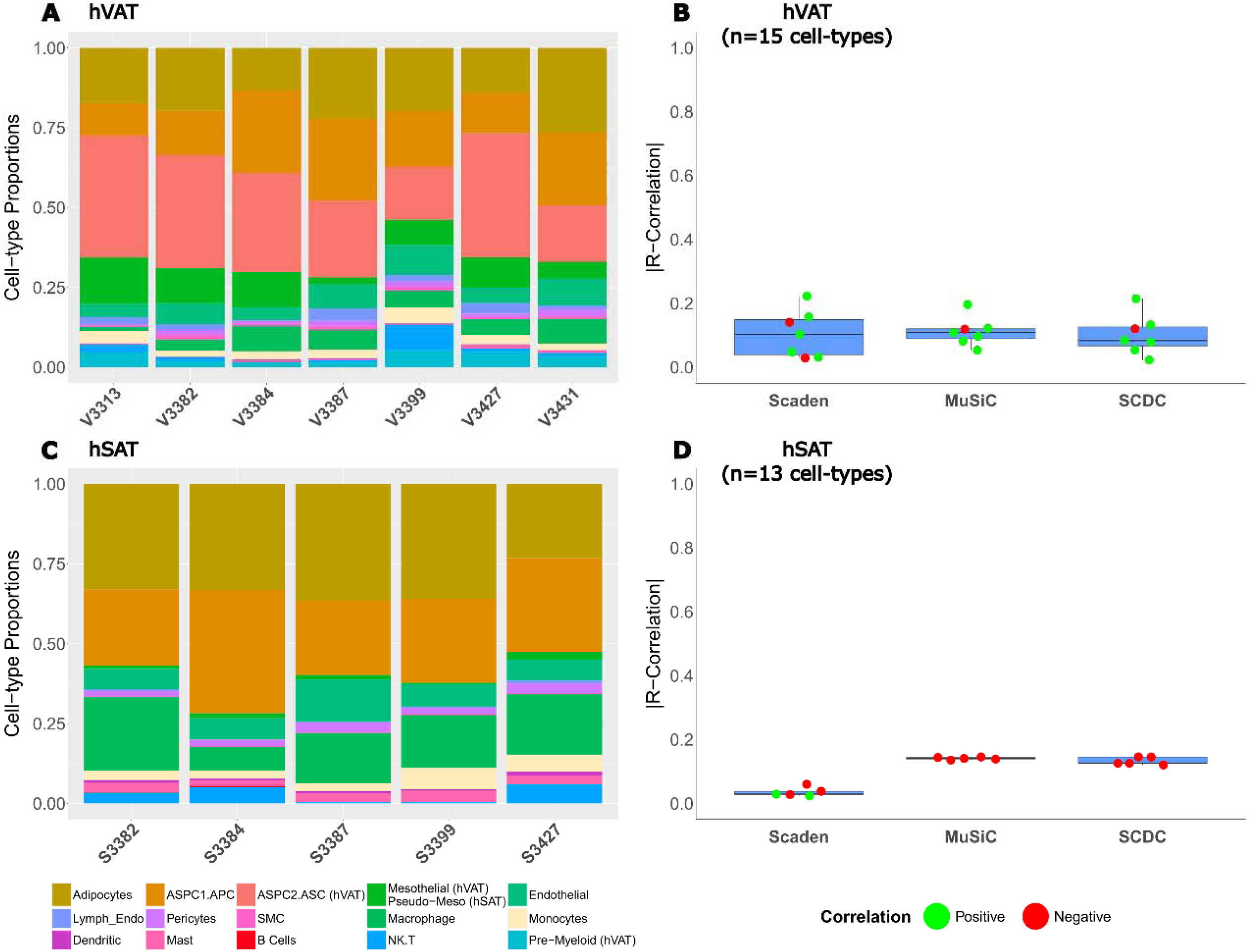
Performance of cell-type estimation by common deconvolution tools analyzing human visceral and subcutaneous adipose tissue bulk RNA-seq data. **A., C.**, Cell-type proportions of 7 hVAT and 5 hSAT samples as assessed by snRNA-seq, respectively. Shown are the proportions of the 15 and 13 cell types, respectively. **B., D.**, Accuracy of estimated cell-type proportions achieved by 3 deconvolution tools: MuSiC and SCDC are deconvolution algorithms that utilize cell-type specific gene expression, and Scaden uses a deep neural network ensemble training approach. Boxplot showing each test-case R correlation coefficient between the true-proportions (snRNA-seq results) and the tools’ estimated proportions for every sample (using leave-one-out training methodology, as detailed in Methods)

The recent human adipose tissue cell atlas estimated cell-type proportions by deconvolution of 331 subcutaneous adipose tissue bulk RNA-seq samples using snRNA-seq data as reference (10). Yet, the estimate was for five major cell-type merged groups: (i) ‘Vascular cells’ included endothelial, lymphatic endothelial, pericytes, and smooth muscle cells; (ii) ‘Myeloid-immune cells’, included macrophages, monocytes, dendritic cells, mast cells and neutrophils; (iii) ‘Lymphoid-immune cells’, included B cells, NK cells and T cells; (iv) ‘Adipocytes’; and (e) ‘adipose stem and progenitor cells’ (ASPCs). We therefore assessed if the performance of the 3 deconvolution tools could be improved by decreasing the resolution of cell-type clustering. For this, we generated “medium” and “low resolution” cell-type merged groups from the 15 and 13 cell-types identified by our snRNA-seq in hVAT and hSAT, respectively. Medium cell type resolution included 10 and 8 cell-type groups, respectively, and low cell type resolution included 5 cell-type groups, as in (10) + 1 group named “other” (**Figure 4A,B**). Decreasing the number of cell-type groups indeed improved the performance of the deconvolution tools, as indicated by an increased coefficient of correlation. While hVAT did not show improved performance (higher correlation coefficient) in the middle resolution, it did exhibit improved performance in the low resolution (**Figure 4C**). In hSAT, the correlation coefficient increased gradually with attempting to estimate a lower number of cell type groups (**Figure 4D**). Overall, the best performance was achieved when attempting to estimate in hVAT, at low cell-type resolution, using Scaden. Nevertheless, the coefficient of correlation even for this analysis was 0.61.

**Figure 4:**
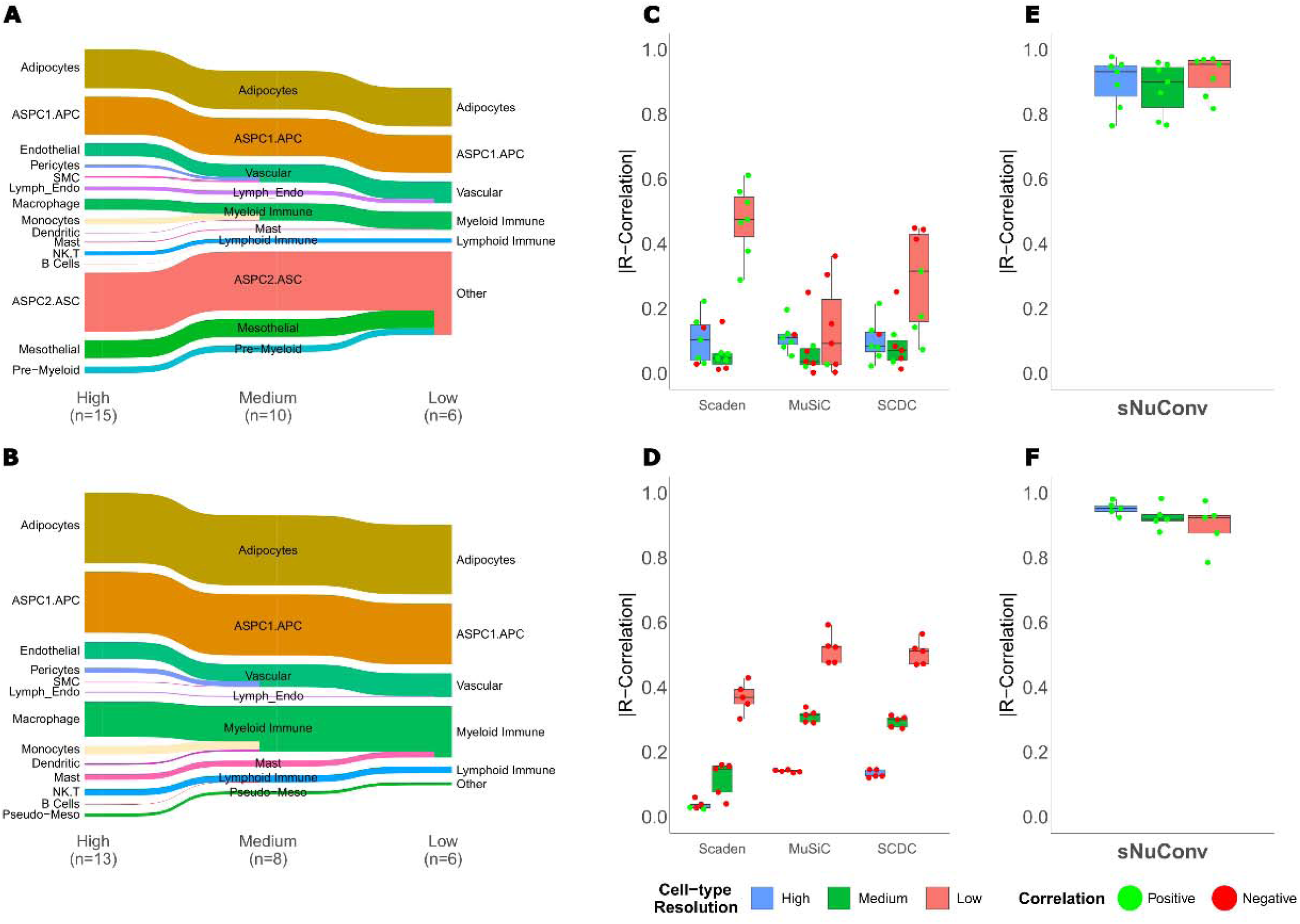
Performance of existing deconvolution algorithms and of sNuConv at various resolutions of cell-types/group. **A., B.**, We merged the “high cell-type resolution” of 15 and 13 specific hVAT and hSAT cell types, respectively, to generate medium and low resolution of cell-type groups, consisting of 10 and 6 cell-type groups, respectively, for hVAT, and 8 and 6, respectively, for hSAT. **C., D.**, The accuracy of cell-type composition estimation at the three cell-type/group resolution levels is shown for Scaden, MuSiC and SCDC, as in Figure 3, for hVAT and hSAT, respectively. **E.**, **F.**, Accuracy of cell-type composition estimation by sNuConv – a Scaden-based deconvolution tool developed to train on snRNA-seq data (as illustrated in Figure 2). Boxplot showing each test-case R correlation coefficient between the true-proportions (snRNA-seq results) and the tools’ estimated proportions for every sample (as in **C.** and **D.**).

Collectively, 3 existing deconvolution algorithms exhibit low reliability in estimating the full set of snRNA-seq -identified cell-types in hVAT and hSAT. They provide a somewhat improved, but still low, capacity to estimate merged cell-type groups. This performance level of existing tools prompted us to develop an improved deconvolution tool for tissues like human adipose tissue, whose true cell-type composition necessitates snRNA-seq, rather than scRNA-seq, analysis.

### 3.2. sNuConv Methodology

We hypothesized that the poor performance of the existing deconvolution tools stems from their reliance on scRNA-seq as reference data for training. Both scRNA-seq (training dataset) and bulk RNA-seq largely represent a common source of the RNA pool, i.e. mature/cytoplasmic mRNA, in contrast to the nuclear RNA pool, which is distinct (12). Additionally, to accurately estimate cell proportions of adipose tissue, the adipocytes must be also considered. This necessitates reliance on snRNA-seq data, and thus – on the nuclear RNA pool, for the algorithms’ training. We aimed to implement a new, snRNA-seq tailored method (sNuConv) that will adjust for these differences in order to better fit the snRNA-seq dataset to the bulk RNA-seq dataset. The fitted (corrected) dataset, which better represents true bulk RNA-seq samples, can then serve for training deep-learning algorithms. We chose to use the deconvolution deep-learning based algorithm Scaden as our core training and prediction method on which we implemented our sNuConv methodology.

#### Per-gene regression model

We took advantage of our paired snRNA-seq and bulk RNA-seq samples from the same donors to bridge the gap between nuclei-based RNA and cytoplasmic mature mRNAs. We set to discover genes whose expression is highly correlated between the nuclei and the cytoplasm (**Figure 2A**_1_). These genes enabled us to transform our snRNA-seq data to become more comparable to single-cell data (**Figure 2A**_2,3_). First, we simulated “pseudo bulk RNA-seq**"** samples based on the snRNA-seq dataset by per-gene averaging of randomly sampled 10,000 cells, repeated 30 times, while reflecting the sample’s true cell-type proportions. The final gene counts for each pseudo-bulk sample were then normalized to Counts per Million (CPM). The specific combination of paired true bulk RNA-seq (also normalized to CPM) and simulated pseudo-bulk samples having true proportions enabled us to look for genes which behave in a correlated manner. A linear regression model was generated for each highly correlated gene (|R|>0.6, Spearman) (**Figure 2A**_1_).

#### Pseudo-bulk Simulation by Scaden using random proportions, gene filtration and correction for sn-bulk correlated genes

In order to create a training dataset that resembles bulk RNA-seq samples, Scaden was used to generate 1,000 simulated pseudo-bulk samples with random (but known) proportions (**Figure 2A**_2_). Each simulation was based on 1,000 cells selected from a pooled snRNA-seq dataset comprised of all samples from the same depot. The gene counts of the 1,000 cells were summed up to create a pseudo-bulk sample. The generated simulated pseudo-bulk samples with random proportions were first normalized to CPM and then filtered to include only the highly correlated genes (from **Figure 2A**_1_). The normalized gene counts were then corrected, each using the specific gene’s regression model calculated from the paired sn-bulk analysis, to yield “corrected pseudo-bulk RNA-seq**"** samples (**Figure 2A**_3_).

#### Deep-Learning training using corrected pseudo-bulk samples, and computing a per cell-type regression model

Using the 1,000 corrected pseudo-bulk RNA-seq dataset, we allowed Scaden to train, while knowing each pseudo-bulk’s (randomly selected) cell-type proportions (**Figure 2B**). Scaden applied its deep learning algorithms (training step) to all **’**pseudo-bulk samples’, and finally generated three prediction models. These models were subsequently applied to estimate cell-type proportions of the true bulk RNA-seq samples (that were also used during the training step) (**Figure 2C**_1_). Next, we calculated the correlation between the true proportions (based on the snRNA-Seq samples) and Scaden’s estimated proportions (**Figure 2C**_2_). We noted that the cell-types differed in the strength of correlation between the true snRNA-seq-based and Scaden’s estimated cell-type proportions. Thus, the model could be further improved by adjusting the final predictions. For this, we generated a cell-type specific regression model (termed "Per-cell type regression model") for each of the cell-types (**Figure 2C**_2_). Cell-types with a calculated R correlation coefficient above a certain cutoff (Default: R^2^>0.8) were used to adjust the predicted proportions of our test bulk RNA-seq data.

#### Cell type proportion estimation on test data with cell-type proportions adjustment

The final cell-type proportion estimation from true bulk RNA-seq data (**Figure 2D**) included: (i) applying the prediction models generated by Scaden on the test bulk RNA-seq sample (which was not part of the training process) to produce estimated cell-type proportions; (ii) Adjusting the proportions using the "Per-cell-type regression model". Due to differences in correlation coefficients, we chose to correct only for the highly correlated cell-types. This ensured that proportions of cell-types with high confidence were estimated more accurately while minimizing the overall estimation error. Consequently, adjustment of certain proportions shifted the total sum of all cell-type proportions from 1. Therefore, shifting was then required for the non-adjusted proportions.

### 3.3. sNuConv outperformed existing deconvolution tools

The performance of sNuConv in estimating cell-type proportions in bulk RNA-seq samples is shown in **Figure 4E, F**. sNuConv achieved median absolute correlations of 0.954, 0.900 and 0.932 in estimating 6, 10, and 15 hVAT cell-types/groups respectively, and 0.924, 0.918 and 0.953 in estimating 6, 8 and 13 cell-types/groups respectively in hSAT. In comparison to the second-highest performing tool in hVAT – Scaden - which exhibited median absolute correlations of 0.474, 0.046 and 0.103 for 6, 10, and 15 hVAT cell-types/groups respectively, sNuConv predictions outperformed Scaden. In hSAT, the second highest performing tool was MuSiC, achieving median absolute correlations of 0.524, 0.315 and 0.141 for 6, 8 and 13 cell-types/groups respectively, again, markedly lower than sNuConv’s performance. Notably, sNuConv was the only tool to estimate cell-types with consistent accuracy, producing only positive correlations in all cell-type/groups for both depots. A similar trend was observed assessing the performance of the different deconvolution tools using Root Mean Square Error (RMSE, data not shown) as a performance accuracy measure.

## 4. Discussion

Deconvolution algorithms have been developed to estimate cell-type composition of tissues from their bulk RNA-seq data. However, existing deconvolution tools largely rely on cell-specific RNA markers derived from single-cell RNA sequencing. Moreover, rarely are the estimations obtained by such deconvolution tools experimentally validated by results of same samples analyzed in parallel by both bulk RNA-seq and snRNA-seq technologies to assess the estimated-to-true correlation. Here, we experimentally analyzed the same samples, from two different adipose tissue depots – hSAT and hVAT – by both methods: bulk and snRNA-seq. Our results demonstrate the following main points:

- Existing deconvolution tools perform poorly in estimating cell-type proportions when trained on snRNA-seq data (instead of scRNA-seq), and applying it to bulk RNA-seq.

- This likely reflects the known differences in the RNA repertoire between the nuclear compartment and the whole-cell. The latter is dominated by cytoplasmic RNA, while nuclear RNA includes larger proportions of not fully processed RNA molecules and long non-coding RNAs (12).

- Thus, we herein show that training a deconvolution algorithm on a subset of genes whose expression patterns correlate well between the snRNA-seq and bulk RNA-seq, along with cell proportion correction, can greatly improve the estimated-to-true correlation coefficient.

- A possible limitation of sNuConv may be in its high tissue-specificity, as the deconvolution algorithm performed best when applied to the same adipose tissue depot it was trained on: A combined model that considered both hSAT and hVAT exhibited lower overall performance in estimating cell types from bulk RNA-seq data (data not shown). However, this finding may reflect both the high sensitivity of the tool to unique expression patterns, and the fact that the seemingly-similar hSAT and hVAT are in fact distinct, as can be observed even at the bulk RNA-seq PCA analysis. A significant strength of sNuConv is its apparent relative independence on the resolution (i.e., number of cell types/groups) it estimates from bulk RNA-seq data.

The cellular landscape of adipose tissue may be tightly linked to obesity complications in cross-sectional analyses. Furthermore, studies suggest that the abundance of specific cell types in adipose tissue can predict response to anti-obesity treatment (24). For example, higher abundance of mast cells in hVAT was shown to predict greater weight-loss response 6 months post-bariatric surgery (25). This proof-of-principle study suggests that information on the cellular composition of adipose tissue could uncover novel predictors of response to treatment, and could thereby help in personalizing anti-obesity therapy. Yet, identifying predictors to therapy requires large cohorts, which is unlikely to be obtainable with snRNA-seq analysis, given its high cost and requirements of still non-standard laboratory equipment. Deconvolution can fill in this missing gap by providing cell-type composition estimates from large cohorts of adipose tissue analyzed by bulk RNA-seq. Our study highlights the need to ensure, experimentally, the robustness and trustworthiness of deconvolution algorithms, before applying their use on large bulk RNA-seq data. Here, this was experimentally achieved by parallel examination of same samples using bulk and snRNA-seq. Importantly, a relatively small number of such samples can already guide the optimization of a validated deconvolution tool – a challenge particularly in need in tissues that require snRNA-seq analysis, such as adipose tissue and the brain.

In summary, we hereby present sNuConv, a bioinformatic approach to adopt deconvolution tools that were designed to perform with scRNA-seq data, to enable extracting cell-type composition of tissues necessitating snRNA-seq.

## List of abbreviations

bulk RNA-seq: bulk RNA sequencing
snRNA-seq: single nucleus RNA sequencing
scRNA-seq: single-cell RNA sequencing
hSAT: human subcutaneous adipose tissue
hVAT: human visceral adipose tissue

## Acknowledgements

This study was supported by grants from the Human Cell Atlas project – the Chan-Zuckerberg initiative, the Deutsche Forschungsgemeinschaft (DFG, German Research Foundation) 209933838: SFB1052: “Obesity mechanisms”, and the Israel Science Foundation (ISF-2176/19).

The authors would like to acknowledge the help of Daphne Perlman in the graphical design of the figures in this article.

