## Supplementary figures and images for "sNucConv: A bulk RNA-seq deconvolution method trained on single-nucleus RNA-seq data to estimate cell-type composition of human subcutaneous and visceral adipose tissues"

### Supplemental Figure 1

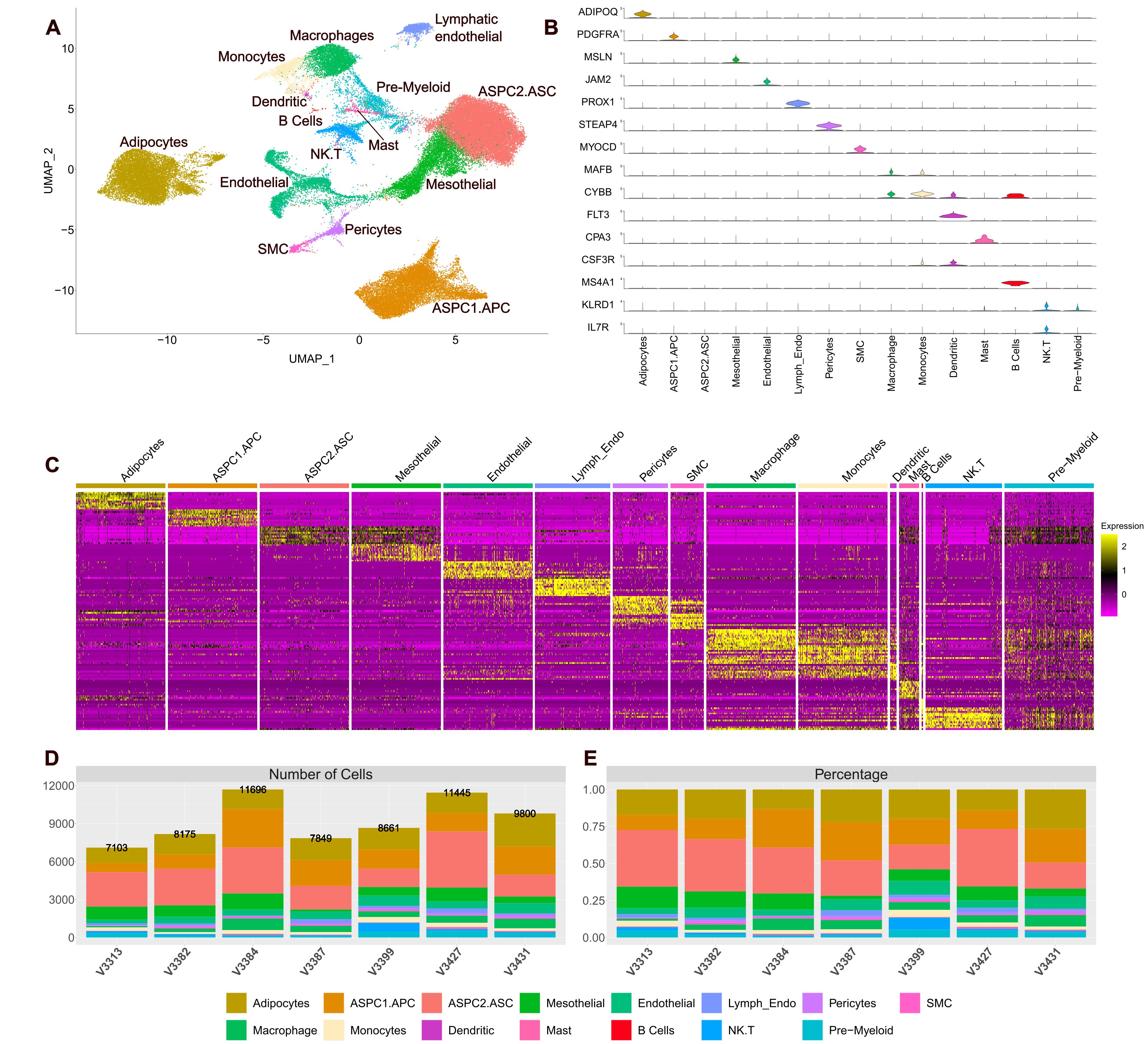

### Supplemental Figure 2

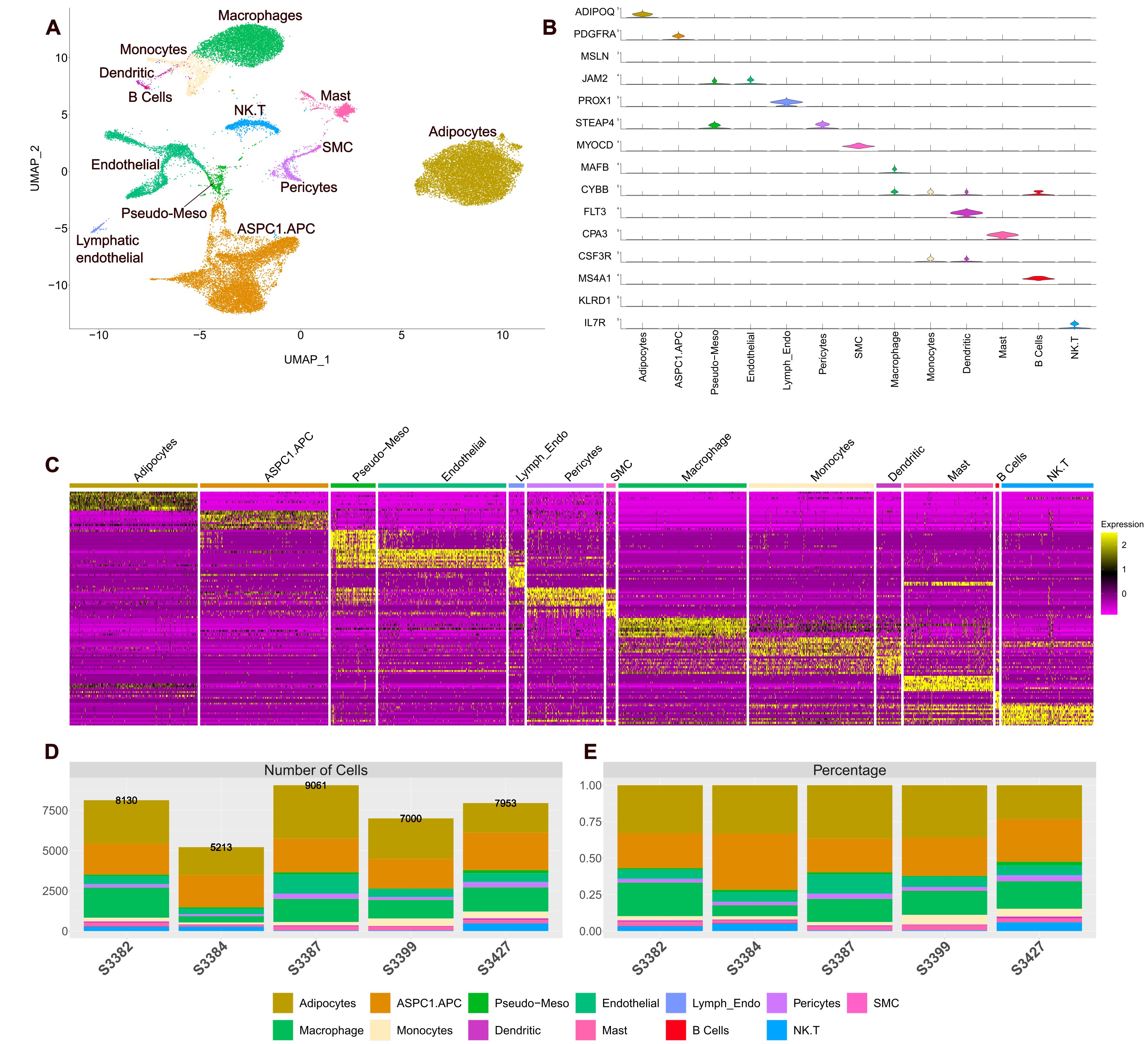
