## Supplemental Table 1 for "sNucConv: A bulk RNA-seq deconvolution method trained on single-nucleus RNA-seq data to estimate cell-type composition of human subcutaneous and visceral adipose tissues"

| **Table 1:** Clinical and Biochemical Characteristics (N=7) | | | | | | | | | | | | |
| --- | --- | --- | --- | --- | --- | --- | --- | --- | --- | --- | --- | --- |
| **Patient number** | | | 1 | | | **2** | **3** | **4** | **5** | **6** | **7** | **Mean** |
|  | | |  | | |  |  |  |  |  |  | *(min-max)* |
| Age (y) | | | 39 | | | 42 | 34 | 34 | 33 | 24 | 49 | **36.4**  *(24.0-49.0)* |
| Sex (male / female) | | | male | | | female | male | female | female | male | female | **3 / 4** |
| Body mass index  (BMI, kg/m^2^) | | | 46.0 | | | 41.0 | 34.5 | 40.9 | 36.2 | 44.1 | 35.7 | **39.8**  *(34.5-46.0)* |
| Diastolic blood pressure (mm Hg) | | | 79.0 | | | 87.0 | 61.0 | 80.0 | 88.0 | 95.0 | 70.0 | **80.0**  *(61.0-95.0)* |
| Systolic blood pressure (mm Hg) | | | 130.0 | | | 123.0 | 116.0 | 117.0 | 133.0 | 132.0 | 128.0 | **125.6**  *(116.0-133.0)* |
| Triglycerides (mg/dL) | | | 114.0 | | | 131.0 | 181.0 | 69.0 | 179.0 | 65.0 | 195.0 | **133.4**  *(65.0-195.0)* |
| Total cholesterol (mg/dL) | | | 101.0 | | | 121.0 | 133.0 | 154.0 | 64.0 | 168.0 | 143.0 | **126.3**  *(64.0-168.0)* |
| Low-density lipoproteins (LDL, mg/dL) | | | 56.0 | | | 89.0 | 119.0 | 99.0 | 28.0 | 102.0 | 56.0 | **78.4**  *(28-119)* |
| high-density lipoproteins (HDL, mg/dL) | | | 38.0 | | |  | 48.0 | 42.0 | 38.0 | 54.0 | 48.0 | **44.7**  *(38.0-54.0)* |
| Fasting plasma glucose (FPG, mg/dL) | | | 124.0 | | | 86.0 | 78.0 | 103.0 | 135.0 | 90.0 | 106.0 | **103.1**  *(78.0-135.0)* |
| Hemoglobin A1c  (HbA_1c_ , %) | | | 5.4 | | | 5.4 | 5.5 | 5.8 | 6.2 | 5.8 | 6.7 | **5.8**  *(5.4-6.7)* |
| C-reactive protein (CRP) | | | 1.47 | | | 0.46 | 0.64 | 0.97 | 0.38 | 0.45 | 0.90 | **0.75** |
|  | | |  | | |  |  |  |  |  |  | *(0.38-1.47)* |
