## Supplemental Table 2 for "sNucConv: A bulk RNA-seq deconvolution method trained on single-nucleus RNA-seq data to estimate cell-type composition of human subcutaneous and visceral adipose tissues"

| **Table 2:** Cell Ranger parameters (raw counts, before QC) | | | | | | | | | | |
| --- | --- | --- | --- | --- | --- | --- | --- | --- | --- | --- |
|  | | **Adipose tissue depot** | | **Estimated number of nuclei** | | | **Mean reads per nucleus** | **Median genes per nucleus** | **Fraction reads in nuclei** | **Sequencing saturation** |
| **1** | | vis | | 11,919 | | | 26,175 | 1,841 | 80.1% | 54.7% |
| **2** | | sc | | 11,215 | | | 54,243 | 1,960 | 53.5% | 61.4% |
|  | | vis | | 9,482 | | | 35,831 | 1,341 | 60.9% | 74.0% |
| **3** | | sc | | 6,860 | | | 88,421 | 1,704 | 50.1% | 76.2% |
| **4**  **5**  **6**  **7** | | vis  sc  vis  sc  vis  sc  vis  vis | | 14,595  10,059  11,419  8,563  11,796  9,203  14,605  12,043 | | | 32,624  45,335  45,930  41,334  34,034  56,577  36,866  41,091 | 2,077  2,613  2,299  2,154  1,822  1,632  1,626  1,362 | 58.6%  70.1%  68.8%  69.7%  68.3%  60.4%  59.2%  59.5% | 32.2%  48.0%  44.1%  40.5%  48.5%  65.2%  56.7%  70.9% |
| **Mean** | | **sc** | | **9,180** | | | **57,182** | **2,020** | **61%** | **58%** |
|  | | **vis** | | **12,265** | | | **36,078** | **1,766** | **65%** | **54%** |
| *min-max* | | *sc* | | *6,860-11,215* | | | *41,334-88,421* | *1,632-2,613* | *50.1%-*70.1% | *40.5%-76.2%* |
|  | | *vis* | | *9,482-14,595* | | | *26,175-45,930* | *1,341-2,299* | *58.6%-80.1%* | *32.2%-74.0%* |
