## Supplemental Table 3 for "sNucConv: A bulk RNA-seq deconvolution method trained on single-nucleus RNA-seq data to estimate cell-type composition of human subcutaneous and visceral adipose tissues"

|  | **Table 3:** Top 10 highest preferentially expressed genes per cluster in hVAT snRNA-seq | | | | | | | | | | | |
| --- | --- | --- | --- | --- | --- | --- | --- | --- | --- | --- | --- | --- |
|  | | | **avg_log2FC** | | **pct.1** | | | | **pct.2** | **p_val_adj** | **cluster** | **gene** |

| 1 | 4.963877 | 0.974 | 0.01 | 0 | Adipocytes | GPAM |
| --- | --- | --- | --- | --- | --- | --- |
| 2 | 4.553834 | 0.984 | 0.131 | 0 | Adipocytes | ACACB |
| 3 | 4.51975 | 0.975 | 0.052 | 0 | Adipocytes | PDE3B |
| 4 | 4.045003 | 0.96 | 0.049 | 0 | Adipocytes | SORBS1 |
| 5 | 3.79457 | 0.709 | 0.045 | 0 | Adipocytes | CLSTN2 |
| 6 | 3.58954 | 0.857 | 0.015 | 0 | Adipocytes | KCNIP2-AS1 |
| 7 | 3.427984 | 0.606 | 0.046 | 0 | Adipocytes | LRP1B |
| 8 | 3.370068 | 0.793 | 0.006 | 0 | Adipocytes | AL136119.1 |
| 9 | 3.360463 | 0.913 | 0.078 | 0 | Adipocytes | DIRC3 |
| 10 | 3.27022 | 0.9 | 0.004 | 0 | Adipocytes | PLIN1 |
| 11 | 4.343157 | 0.933 | 0.172 | 0 | ASPC1.APC | NEGR1 |
| 12 | 3.680945 | 0.859 | 0.176 | 0 | ASPC1.APC | ABCA10 |
| 13 | 3.65085 | 0.866 | 0.209 | 0 | ASPC1.APC | LAMA2 |
| 14 | 3.57427 | 0.761 | 0.036 | 0 | ASPC1.APC | DCN |
| 15 | 3.524017 | 0.632 | 0.051 | 0 | ASPC1.APC | C7 |
| 16 | 3.340494 | 0.35 | 0.069 | 0 | ASPC1.APC | AL590807.1 |
| 17 | 3.21994 | 0.826 | 0.173 | 0 | ASPC1.APC | ABCA9 |
| 18 | 3.214771 | 0.619 | 0.076 | 0 | ASPC1.APC | FGF14 |
| 19 | 3.181134 | 0.683 | 0.144 | 0 | ASPC1.APC | ABCA9-AS1 |
| 20 | 3.069083 | 0.714 | 0.107 | 0 | ASPC1.APC | DCLK1 |
| 21 | 2.316212 | 0.904 | 0.21 | 0 | ASPC2.ASC | AP000561.1 |
| 22 | 2.242101 | 0.76 | 0.178 | 0 | ASPC2.ASC | KCTD8 |
| 23 | 2.222824 | 0.957 | 0.226 | 0 | ASPC2.ASC | PKHD1L1 |
| 24 | 2.151194 | 0.688 | 0.159 | 0 | ASPC2.ASC | RBFOX1 |
| 25 | 2.124657 | 0.869 | 0.244 | 0 | ASPC2.ASC | FAM155A |
| 26 | 2.109198 | 0.731 | 0.159 | 0 | ASPC2.ASC | LINC02360 |
| 27 | 2.08058 | 0.72 | 0.169 | 0 | ASPC2.ASC | AC005699.1 |
| 28 | 2.025913 | 0.808 | 0.247 | 0 | ASPC2.ASC | PTPRQ |
| 29 | 2.02288 | 0.834 | 0.221 | 0 | ASPC2.ASC | THSD4 |
| 30 | 2.000468 | 0.901 | 0.365 | 0 | ASPC2.ASC | SOX6 |
| 31 | 3.11013 | 0.567 | 0.061 | 0 | Mesothelial | ITLN1 |
| 32 | 3.052253 | 0.574 | 0.033 | 0 | Mesothelial | TMSB4X |
| 33 | 2.824265 | 0.554 | 0.034 | 0 | Mesothelial | EEF1A1 |
| 34 | 2.779006 | 0.5 | 0.023 | 0 | Mesothelial | TIMP1 |
| 35 | 2.737877 | 0.482 | 0.02 | 0 | Mesothelial | RPLP1 |
| 36 | 2.679268 | 0.498 | 0.02 | 0 | Mesothelial | RPL41 |
| 37 | 2.608321 | 0.457 | 0.018 | 0 | Mesothelial | RPL13 |
| 38 | 2.557908 | 0.633 | 0.055 | 0 | Mesothelial | RPS21 |
| 39 | 2.544715 | 0.529 | 0.061 | 0 | Mesothelial | MT-CO3 |
| 40 | 2.536788 | 0.451 | 0.017 | 0 | Mesothelial | RPL3 |
| 41 | 4.727634 | 0.872 | 0.033 | 0 | Endothelial | MECOM |
| 42 | 3.584102 | 0.716 | 0.02 | 0 | Endothelial | ANO2 |
| 43 | 3.570551 | 0.733 | 0.02 | 0 | Endothelial | VWF |
| 44 | 3.481678 | 0.514 | 0.046 | 0 | Endothelial | BTNL9 |
| 45 | 3.478512 | 0.611 | 0.038 | 0 | Endothelial | PIK3R3 |
| 46 | 3.476022 | 0.692 | 0.016 | 0 | Endothelial | ADGRL4 |
| 47 | 3.434312 | 0.72 | 0.026 | 0 | Endothelial | EMCN |
| 48 | 3.305172 | 0.624 | 0.013 | 0 | Endothelial | FLT1 |
| 49 | 3.296854 | 0.736 | 0.031 | 0 | Endothelial | PTPRB |
| 50 | 3.242993 | 0.786 | 0.426 | 0 | Endothelial | ARL15 |
| 51 | 4.389819 | 0.87 | 0.009 | 0 | Lymph_Endo | MMRN1 |
| 52 | 4.179641 | 0.799 | 0.016 | 0 | Lymph_Endo | RELN |
| 53 | 4.021756 | 0.845 | 0.033 | 0 | Lymph_Endo | LINC02147 |
| 54 | 3.993559 | 0.583 | 0.03 | 0 | Lymph_Endo | AL357507.1 |
| 55 | 3.846931 | 0.686 | 0.025 | 0 | Lymph_Endo | LINC02208 |
| 56 | 3.774554 | 0.66 | 0.014 | 0 | Lymph_Endo | NRG3 |
| 57 | 3.766171 | 0.931 | 0.163 | 0 | Lymph_Endo | TFPI |
| 58 | 3.668894 | 0.892 | 0.093 | 0 | Lymph_Endo | STOX2 |
| 59 | 3.43866 | 0.724 | 0.012 | 0 | Lymph_Endo | PROX1 |
| 60 | 3.234556 | 0.783 | 0.091 | 0 | Lymph_Endo | SNTG2 |
| 61 | 5.142308 | 0.889 | 0.046 | 0 | Pericytes | COL25A1 |
| 62 | 3.575982 | 0.715 | 0.073 | 0 | Pericytes | PDE1C |
| 63 | 3.404479 | 0.724 | 0.081 | 0 | Pericytes | ADGRB3 |
| 64 | 3.390147 | 0.781 | 0.168 | 0 | Pericytes | RGS6 |
| 65 | 3.30472 | 0.765 | 0.129 | 0 | Pericytes | INPP4B |
| 66 | 3.303867 | 0.695 | 0.036 | 0 | Pericytes | STEAP4 |
| 67 | 3.292744 | 0.684 | 0.022 | 0 | Pericytes | LINC00989 |
| 68 | 3.256497 | 0.566 | 0.054 | 0 | Pericytes | NCKAP5 |
| 69 | 3.22986 | 0.952 | 0.519 | 0 | Pericytes | DLC1 |
| 70 | 3.144809 | 0.766 | 0.151 | 0 | Pericytes | LINC02237 |
| 71 | 4.306931 | 0.911 | 0.057 | 0 | SMC | RCAN2 |
| 72 | 3.976717 | 0.851 | 0.059 | 0 | SMC | MYH11 |
| 73 | 3.748526 | 0.936 | 0.17 | 0 | SMC | RGS6 |
| 74 | 3.664793 | 0.491 | 0.036 | 0 | SMC | ADGRL3 |
| 75 | 3.469526 | 0.911 | 0.178 | 0 | SMC | CARMN |
| 76 | 3.066429 | 0.911 | 0.244 | 0 | SMC | CACNA1C |
| 77 | 2.973184 | 0.555 | 0.016 | 0 | SMC | DGKB |
| 78 | 2.926589 | 0.861 | 0.321 | 0 | SMC | ZFHX3 |
| 79 | 2.902464 | 0.632 | 0.038 | 0 | SMC | NTRK3 |
| 80 | 2.876875 | 0.626 | 0.018 | 0 | SMC | DGKG |
| 81 | 4.479829 | 0.9 | 0.058 | 0 | Macrophage | F13A1 |
| 82 | 4.138062 | 0.85 | 0.051 | 0 | Macrophage | LGMN |
| 83 | 4.066435 | 0.901 | 0.063 | 0 | Macrophage | MRC1 |
| 84 | 3.518582 | 0.718 | 0.056 | 0 | Macrophage | SCN1A-AS1 |
| 85 | 3.38028 | 0.736 | 0.037 | 0 | Macrophage | CD163L1 |
| 86 | 3.348881 | 0.829 | 0.076 | 0 | Macrophage | IQGAP2 |
| 87 | 3.320504 | 0.781 | 0.047 | 0 | Macrophage | MS4A4A |
| 88 | 3.302796 | 0.879 | 0.274 | 0 | Macrophage | MTSS1 |
| 89 | 3.283608 | 0.77 | 0.19 | 0 | Macrophage | AFF3 |
| 90 | 3.252223 | 0.955 | 0.377 | 0 | Macrophage | SLC9A9 |
| 91 | 3.103542 | 0.883 | 0.327 | 0 | Monocytes | RBPJ |
| 92 | 2.956435 | 0.901 | 0.21 | 0 | Monocytes | RBM47 |
| 93 | 2.906604 | 0.571 | 0.011 | 0 | Monocytes | MARCO |
| 94 | 2.875353 | 0.767 | 0.063 | 0 | Monocytes | CTSB |
| 95 | 2.802527 | 0.406 | 0.1 | 0 | Monocytes | FN1 |
| 96 | 2.646704 | 0.709 | 0.04 | 0 | Monocytes | CYBB |
| 97 | 2.625009 | 0.757 | 0.066 | 0 | Monocytes | FYB1 |
| 98 | 2.606676 | 0.728 | 0.063 | 0 | Monocytes | CD163 |
| 99 | 2.602877 | 0.663 | 0.143 | 0 | Monocytes | KCNMA1 |
| 100 | 2.593009 | 0.803 | 0.166 | 0 | Monocytes | BMP2K |
| 101 | 3.833325 | 1 | 0.006 | 1.43E-304 | Dendritic | FLT3 |
| 102 | 3.299205 | 0.795 | 0.058 | 1.71E-123 | Dendritic | CIITA |
| 103 | 4.049917 | 0.67 | 0.043 | 6.81E-116 | Dendritic | WDFY4 |
| 104 | 2.978479 | 0.759 | 0.037 | 8.09E-104 | Dendritic | HLA-DPB1 |
| 105 | 3.441021 | 0.902 | 0.24 | 2.12E-101 | Dendritic | HDAC9 |
| 106 | 3.070229 | 0.857 | 0.08 | 1.52E-91 | Dendritic | CD74 |
| 107 | 2.9604 | 0.786 | 0.206 | 7.52E-91 | Dendritic | CPVL |
| 108 | 3.032515 | 0.786 | 0.088 | 1.32E-79 | Dendritic | AOAH |
| 109 | 3.261764 | 0.75 | 0.227 | 3.31E-67 | Dendritic | CCSER1 |
| 110 | 3.618542 | 0.268 | 0.005 | 2.20E-66 | Dendritic | CLNK |
| 111 | 3.338115 | 0.849 | 0.006 | 0 | Mast | KIT |
| 112 | 2.940266 | 0.66 | 0.003 | 0 | Mast | CPA3 |
| 113 | 2.854465 | 0.604 | 0.004 | 0 | Mast | AC092979.1 |
| 114 | 2.471298 | 0.598 | 0.007 | 0 | Mast | HPGD |
| 115 | 2.538252 | 0.663 | 0.026 | 1.07E-269 | Mast | IL18R1 |
| 116 | 2.307968 | 0.618 | 0.022 | 1.08E-253 | Mast | SLC18A2 |
| 117 | 2.494351 | 0.725 | 0.048 | 1.81E-238 | Mast | PZP |
| 118 | 2.272984 | 0.592 | 0.026 | 3.60E-218 | Mast | HPGDS |
| 119 | 2.602124 | 0.772 | 0.114 | 5.75E-212 | Mast | SYTL3 |
| 120 | 2.312108 | 0.95 | 0.427 | 1.60E-172 | Mast | NTM |
| 121 | 4.64485 | 0.913 | 0.113 | 1.65E-76 | B Cells | BANK1 |
| 122 | 3.330398 | 1 | 0.001 | 1.25E-67 | B Cells | MS4A1 |
| 123 | 3.165012 | 0.739 | 0.002 | 1.05E-39 | B Cells | PAX5 |
| 124 | 3.669437 | 0.696 | 0.051 | 3.88E-36 | B Cells | AP002075.1 |
| 125 | 3.37476 | 0.696 | 0.014 | 4.34E-32 | B Cells | COL19A1 |
| 126 | 3.70006 | 0.913 | 0.117 | 5.66E-26 | B Cells | PRKCB |
| 127 | 3.164494 | 0.957 | 0.124 | 2.06E-22 | B Cells | RIPOR2 |
| 128 | 3.506135 | 0.826 | 0.24 | 2.35E-22 | B Cells | BACH2 |
| 129 | 3.518697 | 0.957 | 0.222 | 1.30E-21 | B Cells | AFF3 |
| 130 | 3.131971 | 1 | 0.146 | 2.47E-20 | B Cells | ARHGAP15 |
| 131 | 4.274297 | 0.885 | 0.033 | 0 | NK.T | SKAP1 |
| 132 | 4.018835 | 0.795 | 0.106 | 0 | NK.T | AC079793.1 |
| 133 | 3.729154 | 0.892 | 0.131 | 0 | NK.T | ARHGAP15 |
| 134 | 3.686006 | 0.668 | 0.019 | 0 | NK.T | THEMIS |
| 135 | 3.597314 | 0.895 | 0.101 | 0 | NK.T | PTPRC |
| 136 | 3.259106 | 0.621 | 0.023 | 0 | NK.T | CD247 |
| 137 | 3.165587 | 0.624 | 0.018 | 0 | NK.T | BCL11B |
| 138 | 3.135934 | 0.679 | 0.114 | 0 | NK.T | RIPOR2 |
| 139 | 3.07476 | 0.611 | 0.023 | 0 | NK.T | SAMD3 |
| 140 | 2.992194995 | 0.699 | 0.069 | 0 | NK.T | IKZF1 |
| 141 | 1.032045 | 0.51 | 0.081 | 0 | Pre-Myeloid | LGMN |
| 142 | 0.951684 | 0.408 | 0.038 | 0 | Pre-Myeloid | SKAP1 |
| 143 | 0.912575 | 0.637 | 0.1 | 0 | Pre-Myeloid | IQGAP2 |
| 144 | 0.907002 | 0.612 | 0.1 | 0 | Pre-Myeloid | PTPRC |
| 145 | 0.865534 | 0.536 | 0.095 | 0 | Pre-Myeloid | MRC1 |
| 146 | 0.854247 | 0.534 | 0.09 | 0 | Pre-Myeloid | F13A1 |
| 147 | 0.84324 | 0.425 | 0.081 | 0 | Pre-Myeloid | SCN1A-AS1 |
| 148 | 0.832859 | 0.604 | 0.131 | 0 | Pre-Myeloid | ARHGAP15 |
| 149 | 0.811092 | 0.871 | 0.371 | 0 | Pre-Myeloid | FRMD4B |
| 150 | 0.808878 | 0.509 | 0.107 | 0 | Pre-Myeloid | AC079793.1 |
