## Supplemental Table 4 for "sNucConv: A bulk RNA-seq deconvolution method trained on single-nucleus RNA-seq data to estimate cell-type composition of human subcutaneous and visceral adipose tissues"

|  | **Table 4:** Top 10 highest preferentially expressed genes per cluster in hSAT snRNA-seq | | | | | | | | | | | |
| --- | --- | --- | --- | --- | --- | --- | --- | --- | --- | --- | --- | --- |
|  | | | **avg_log2FC** | | **pct.1** | | | | **pct.2** | **p_val_adj** | **cluster** | **gene** |

| 1 | 5.177445886 | 0.984 | 0.104 | 0 | Adipocytes | WDPCP |
| --- | --- | --- | --- | --- | --- | --- |
| 2 | 4.237290241 | 0.922 | 0.003 | 0 | Adipocytes | GPAM |
| 3 | 3.817781561 | 0.931 | 0.054 | 0 | Adipocytes | CAV2 |
| 4 | 3.793963006 | 0.978 | 0.073 | 0 | Adipocytes | PDE3B |
| 5 | 3.770687236 | 0.879 | 0.025 | 0 | Adipocytes | CLSTN2 |
| 6 | 3.600163042 | 0.958 | 0.09 | 0 | Adipocytes | PCDH9 |
| 7 | 3.592146869 | 0.972 | 0.051 | 0 | Adipocytes | SORBS1 |
| 8 | 3.447522812 | 0.959 | 0.001 | 0 | Adipocytes | PLIN1 |
| 9 | 3.442499973 | 0.87 | 0.007 | 0 | Adipocytes | KCNIP2-AS1 |
| 10 | 3.299245281 | 0.956 | 0.078 | 0 | Adipocytes | LIPE-AS1 |
| 11 | 4.091478426 | 0.93 | 0.148 | 0 | ASPC1.APC | NEGR1 |
| 12 | 4.043650443 | 0.924 | 0.079 | 0 | ASPC1.APC | DCLK1 |
| 13 | 4.026382542 | 0.628 | 0.008 | 0 | ASPC1.APC | DCN |
| 14 | 3.575108733 | 0.804 | 0.025 | 0 | ASPC1.APC | COL1A2 |
| 15 | 3.507291717 | 0.812 | 0.084 | 0 | ASPC1.APC | BICC1 |
| 16 | 3.482039766 | 0.698 | 0.061 | 0 | ASPC1.APC | LINC02511 |
| 17 | 3.473313314 | 0.391 | 0.031 | 0 | ASPC1.APC | PDZRN4 |
| 18 | 3.388619934 | 0.924 | 0.383 | 0 | ASPC1.APC | NOVA1 |
| 19 | 3.373411574 | 0.654 | 0.063 | 0 | ASPC1.APC | ROBO2 |
| 20 | 3.348642199 | 0.873 | 0.298 | 0 | ASPC1.APC | LAMA2 |
| 21 | 4.444231207 | 0.896 | 0.024 | 0 | Endothelial | MECOM |
| 22 | 3.708634085 | 0.949 | 0.207 | 0 | Endothelial | LDB2 |
| 23 | 3.646991148 | 0.76 | 0.023 | 0 | Endothelial | ADGRL4 |
| 24 | 3.622144111 | 0.766 | 0.02 | 0 | Endothelial | ANO2 |
| 25 | 3.430162968 | 0.524 | 0.037 | 0 | Endothelial | BTNL9 |
| 26 | 3.403997882 | 0.777 | 0.021 | 0 | Endothelial | VWF |
| 27 | 3.378355237 | 0.354 | 0.074 | 0 | Endothelial | CADM2 |
| 28 | 3.278317286 | 0.66 | 0.025 | 0 | Endothelial | PLCB4 |
| 29 | 3.269268948 | 0.701 | 0.276 | 0 | Endothelial | KIAA1217 |
| 30 | 3.269130688 | 0.72 | 0.059 | 0 | Endothelial | ST6GALNAC3 |
| 31 | 4.224304751 | 0.813 | 0.014 | 0 | Lymph_Endo | PKHD1L1 |
| 32 | 3.388735229 | 0.753 | 0.008 | 1.35E-265 | Lymph_Endo | PROX1 |
| 33 | 3.96790601 | 0.775 | 0.008 | 3.18E-264 | Lymph_Endo | MMRN1 |
| 34 | 3.359620473 | 0.835 | 0.222 | 9.86E-214 | Lymph_Endo | TFPI |
| 35 | 3.853912022 | 0.769 | 0.015 | 8.34E-204 | Lymph_Endo | NRG3 |
| 36 | 3.408254178 | 0.769 | 0.05 | 2.83E-181 | Lymph_Endo | TSPAN5 |
| 37 | 3.481882018 | 0.648 | 0.027 | 6.62E-177 | Lymph_Endo | RELN |
| 38 | 3.699628213 | 0.791 | 0.034 | 6.22E-172 | Lymph_Endo | LINC02147 |
| 39 | 3.940272594 | 0.758 | 0.032 | 6.64E-161 | Lymph_Endo | LINC02208 |
| 40 | 3.900862094 | 0.522 | 0.038 | 2.52E-148 | Lymph_Endo | AL357507.1 |
| 41 | 4.41048404 | 0.73 | 0.072 | 0 | Pericytes | COL25A1 |
| 42 | 4.024002123 | 0.843 | 0.079 | 0 | Pericytes | RGS6 |
| 43 | 3.573944358 | 0.977 | 0.266 | 0 | Pericytes | PRKG1 |
| 44 | 3.330708062 | 0.709 | 0.074 | 0 | Pericytes | MYO1B |
| 45 | 3.322367163 | 0.646 | 0.075 | 0 | Pericytes | NCKAP5 |
| 46 | 3.02724831 | 0.758 | 0.134 | 0 | Pericytes | EBF2 |
| 47 | 2.873619467 | 0.446 | 0.022 | 0 | Pericytes | POSTN |
| 48 | 2.850959878 | 0.803 | 0.216 | 0 | Pericytes | AC012409.2 |
| 49 | 2.720470128 | 0.768 | 0.191 | 0 | Pericytes | NR2F2-AS1 |
| 50 | 3.217077701 | 0.342 | 0.036 | 3.65E-291 | Pericytes | ADGRL3 |
| 51 | 3.121940143 | 1 | 0.004 | 5.17E-261 | SMC | MYOCD |
| 52 | 3.44811399 | 0.809 | 0.018 | 1.06E-145 | SMC | RYR2 |
| 53 | 4.309562368 | 0.855 | 0.041 | 2.13E-136 | SMC | ADGRL3 |
| 54 | 3.688122799 | 0.891 | 0.087 | 4.04E-121 | SMC | RCAN2 |
| 55 | 3.630150434 | 0.836 | 0.085 | 4.87E-119 | SMC | SORBS2 |
| 56 | 3.510462378 | 0.791 | 0.091 | 2.32E-118 | SMC | MYH11 |
| 57 | 3.364970764 | 0.991 | 0.281 | 7.59E-112 | SMC | PRKG1 |
| 58 | 3.652452383 | 0.9 | 0.095 | 3.24E-102 | SMC | RGS6 |
| 59 | 3.057268892 | 0.891 | 0.229 | 9.07E-91 | SMC | AC012409.2 |
| 60 | 3.221272592 | 0.427 | 0.073 | 2.99E-52 | SMC | DEC1 |
| 61 | 4.649439376 | 0.953 | 0.047 | 0 | Macrophage | F13A1 |
| 62 | 4.139491032 | 0.905 | 0.044 | 0 | Macrophage | LGMN |
| 63 | 4.028774797 | 0.881 | 0.06 | 0 | Macrophage | SCN1A-AS1 |
| 64 | 3.76873671 | 0.979 | 0.133 | 0 | Macrophage | FRMD4B |
| 65 | 3.667077325 | 0.916 | 0.047 | 0 | Macrophage | MRC1 |
| 66 | 3.642552202 | 0.929 | 0.168 | 0 | Macrophage | PDE4D |
| 67 | 3.574243219 | 0.936 | 0.343 | 0 | Macrophage | RBPJ |
| 68 | 3.43310946 | 0.849 | 0.033 | 0 | Macrophage | SCN9A |
| 69 | 3.289899232 | 0.889 | 0.241 | 0 | Macrophage | NAV2 |
| 70 | 3.084925213 | 0.749 | 0.031 | 0 | Macrophage | CD163L1 |
| 71 | 3.349251032 | 0.611 | 0.087 | 0 | Monocytes | ALCAM |
| 72 | 2.631433586 | 0.466 | 0.014 | 0 | Monocytes | CDCP1 |
| 73 | 2.50999042 | 0.653 | 0.096 | 0 | Monocytes | CPB2-AS1 |
| 74 | 2.483514556 | 0.685 | 0.126 | 0 | Monocytes | AOAH |
| 75 | 2.323708186 | 0.808 | 0.222 | 0 | Monocytes | CHST11 |
| 76 | 2.276033311 | 0.42 | 0.075 | 0 | Monocytes | DOCK3 |
| 77 | 2.266188985 | 0.849 | 0.259 | 0 | Monocytes | SLC8A1 |
| 78 | 2.242567456 | 0.72 | 0.168 | 0 | Monocytes | LYN |
| 79 | 2.236375997 | 0.546 | 0.022 | 0 | Monocytes | ITGAX |
| 80 | 2.120613552 | 0.591 | 0.192 | 0 | Monocytes | TBC1D8 |
| 81 | 3.029344539 | 0.832 | 0.003 | 0 | Dendritic | FLT3 |
| 82 | 3.796836998 | 0.663 | 0.085 | 1.08E-250 | Dendritic | WDFY4 |
| 83 | 3.114179204 | 0.639 | 0.038 | 3.33E-215 | Dendritic | AC120193.1 |
| 84 | 2.594109919 | 0.742 | 0.095 | 4.53E-192 | Dendritic | CIITA |
| 85 | 3.281794878 | 0.419 | 0.031 | 5.26E-176 | Dendritic | AC060234.3 |
| 86 | 2.564328137 | 0.725 | 0.141 | 6.00E-169 | Dendritic | RAB11FIP1 |
| 87 | 2.595034008 | 0.577 | 0.035 | 1.96E-165 | Dendritic | HLA-DQA1 |
| 88 | 3.172700829 | 0.742 | 0.162 | 1.31E-152 | Dendritic | CCSER1 |
| 89 | 2.614442199 | 0.684 | 0.182 | 6.72E-130 | Dendritic | CPVL |
| 90 | 2.479891941 | 0.309 | 0.022 | 2.50E-106 | Dendritic | CCDC26 |
| 91 | 4.233589861 | 0.879 | 0.006 | 0 | Mast | KIT |
| 92 | 4.166356344 | 0.638 | 0.018 | 0 | Mast | LINC02208 |
| 93 | 4.131008918 | 0.796 | 0.008 | 0 | Mast | RAB27B |
| 94 | 3.974810465 | 0.752 | 0.003 | 0 | Mast | AC092979.1 |
| 95 | 3.911527415 | 0.766 | 0.015 | 0 | Mast | HPGD |
| 96 | 3.908662361 | 0.892 | 0.21 | 0 | Mast | SLC24A3 |
| 97 | 3.739809287 | 0.707 | 0.018 | 0 | Mast | LINC02147 |
| 98 | 3.736037941 | 0.975 | 0.374 | 0 | Mast | NTM |
| 99 | 3.708745195 | 0.879 | 0.132 | 0 | Mast | SYTL3 |
| 100 | 3.689878253 | 0.742 | 0.004 | 0 | Mast | CPA3 |
| 101 | 4.98517868 | 0.952 | 0.165 | 1.33E-208 | B Cells | BANK1 |
| 102 | 3.702922194 | 1 | 0.001 | 2.66E-118 | B Cells | MS4A1 |
| 103 | 4.325459651 | 0.762 | 0.105 | 8.08E-101 | B Cells | AP002075.1 |
| 104 | 4.151109673 | 0.905 | 0.136 | 4.59E-88 | B Cells | RALGPS2 |
| 105 | 4.185462895 | 0.81 | 0.001 | 4.26E-86 | B Cells | FCRL1 |
| 106 | 3.726703172 | 0.714 | 0.002 | 2.31E-72 | B Cells | PAX5 |
| 107 | 4.501864657 | 0.81 | 0.145 | 3.27E-71 | B Cells | BACH2 |
| 108 | 4.232764096 | 0.952 | 0.121 | 1.49E-66 | B Cells | PRKCB |
| 109 | 4.012678471 | 0.952 | 0.062 | 1.64E-51 | B Cells | RIPOR2 |
| 110 | 3.462256582 | 0.619 | 0.042 | 6.77E-32 | B Cells | AC120193.1 |
| 111 | 3.912725812 | 0.862 | 0.05 | 0 | NK.T | SKAP1 |
| 112 | 3.847122995 | 0.805 | 0.041 | 0 | NK.T | RIPOR2 |
| 113 | 3.58376336 | 0.565 | 0.01 | 0 | NK.T | THEMIS |
| 114 | 3.480182957 | 0.886 | 0.203 | 0 | NK.T | AC079793.1 |
| 115 | 3.478609974 | 0.631 | 0.021 | 0 | NK.T | CD247 |
| 116 | 3.468385776 | 0.631 | 0.009 | 0 | NK.T | BCL11B |
| 117 | 3.44009934 | 0.909 | 0.208 | 0 | NK.T | ARHGAP15 |
| 118 | 3.263755046 | 0.589 | 0.077 | 0 | NK.T | INPP4B |
| 119 | 3.118692986 | 0.916 | 0.216 | 0 | NK.T | PTPRC |
| 120 | 3.101473575 | 0.516 | 0.015 | 0 | NK.T | SAMD3 |
| 121 | 3.923305972 | 0.748 | 0.175 | 0 | Pseudo-Meso | FABP4 |
| 122 | 3.82098232 | 0.828 | 0.027 | 0 | Pseudo-Meso | RPLP1 |
| 123 | 3.748229775 | 0.669 | 0.039 | 0 | Pseudo-Meso | B2M |
| 124 | 3.733619196 | 0.746 | 0.016 | 0 | Pseudo-Meso | MT-CO3 |
| 125 | 3.529254795 | 0.66 | 0.024 | 0 | Pseudo-Meso | RPL41 |
| 126 | 3.503744463 | 0.769 | 0.021 | 0 | Pseudo-Meso | MT-CYB |
| 127 | 3.493749155 | 0.583 | 0.024 | 0 | Pseudo-Meso | TMSB4X |
| 128 | 3.47512159 | 0.702 | 0.011 | 0 | Pseudo-Meso | MT-ATP6 |
| 129 | 3.461617396 | 0.746 | 0.045 | 0 | Pseudo-Meso | EEF1A1 |
| 130 | 3.457844366 | 0.763 | 0.021 | 0 | Pseudo-Meso | MT-CO1 |
